# Programmable gene regulation for metabolic engineering using decoy transcription factor binding sites

**DOI:** 10.1101/2020.05.05.079665

**Authors:** Tiebin Wang, Nathan Tague, Stephen Whelan, Mary J. Dunlop

**Affiliations:** Molecular Biology, Cell Biology & Biochemistry, Boston University, Boston, MA, 02215, USA; Biological Design Center, Boston University, Boston, MA, 02215, USA; Biomedical Engineering, Boston University, Boston, MA, 02215, USA; Chemistry, Boston University, Boston, MA, 02215, USA

## Abstract

Transcription factor decoy binding sites are short DNA sequences that can serve as “sponges” to titrate a transcription factor away from its natural binding site, therefore regulating gene expression. In this study, we harness decoy sites to develop synthetic transcription factor sponge systems to regulate gene expression for metabolic pathways in *Escherichia coli.* We show that transcription factor sponges can effectively regulate expression of native and heterologous genes. Tunability of the sponge can be engineered via changes in copy number or modifications to the DNA decoy site sequence. Using arginine biosynthesis as a showcase, we observe a 16-fold increase in arginine production when we introduce the sponge system to steer metabolic flux towards increased arginine biosynthesis, with negligible growth differences compared to the wild type strain. The sponge-based production strain shows high genetic stability; in contrast to a gene knock-out approach where mutations were common, we detected no mutations in the production system using the sponge-based strain. We further show that transcription factor sponges are amenable to multiplexed library screening by demonstrating enhanced tolerance to pinene with a combinatorial sponge library. Our study shows that transcription factor sponges are a powerful and compact tool for metabolic engineering.

## INTRODUCTION

Metabolic flux in microbes is coordinated by transcription factors that dictate gene expression levels for genes encoding enzymes that carry out necessary chemical conversions. Removing the effect of transcription factors can alter expression of genes and redirect native metabolic pathways (1, 2). Strategies for doing this can generally be divided into two groups: complete removal and partial removal of the transcription factor. For the first, traditional gene knock-out strategies can be used to remove the gene encoding the transcription factor. This approach has been shown to enhance microbial tolerance towards biofuels (3–6) and increase production of amino acids (7–11). Although techniques for generating gene knock-out are well-established, genome editing is a labor intensive process and can be difficult to multiplex. In addition, since transcription factors can play broad physiological roles, completely removing the gene is frequently associated with detrimental effects, such as fitness costs (7, 12). For example, knocking out *argR* in *E. coli* can substantially increase the production of arginine, but reduces cell growth by 50% (7). This growth deficit inevitably leads to selective pressure against engineered cells, potentially reducing genetic stability, which is an important consideration during the scale up process (13–17).

Partial removal strategies include knock-down approaches such as using CRISPRi or sRNA to downregulate gene expression (7, 10, 18). These approaches are straightforward to design and can be used to achieve multiplexed regulation. Importantly, partially removing the effect of a transcription factor can redirect metabolic flux at intermediate levels that are better tolerated by the cell (7, 10). For example, compared to the 50% growth deficit resulting from an *argR* knock-out, knock-down of *argR* with CRISPRi in *E. coli* reduced growth by 30% while maintaining similar arginine productivity (7). Such partial removal strategies have been employed to increase yields for various target products, such as the nylon precursor cadaverine in addition to the amino acid arginine (7, 10). However, CRISPRi can show significant toxicity and off-target activity (19, 20). In addition, the size of knock-down systems is large and may place a practical limit on the number of genes that can be introduced into the host cell. For example, CRISPRi knockdown requires dCas9 and the associated RNAs, accounting for approximately 5 kilobases of additional genetic material. As a result, it remains challenging to control transcription factor levels using existing design approaches. The ideal strategy would be straightforward to design and construct, have low fitness cost, and exhibit high compatibility with the large heterologous pathways involved in metabolic engineering.

Decoy binding sites provide a potential platform for regulation of transcription factor activity. Decoy binding sites or transcription factor “sponges” are short DNA sequences that can soak up free transcription factors in the cell, titrating them away from their functional promoters to alter gene expression (21–25). Transcription factor sponge sequences are short (~30bp), their strength can be tuned through copy-number or sequence modifications, and they do not rely on transcription or translation activity. Their design and construction are straightforward and are highly amenable to multiplexed regulation. So far, the application of transcription factor sponges in synthetic biology has been limited to synthetic gene circuits to either alter dynamics or lower noise (25, 26). However, the application of transcription factor sponges to the regulation of endogenous genes and metabolic pathways remains largely unexplored. Here, we seek to demonstrate the utility of transcription factor sponges for metabolic engineering.

We designed and engineered sponges that allow multiplexed and tunable regulation of transcription factors related to metabolism. We show that the synthetic sponge systems can effectively alter transcriptional outputs from downstream promoters of targeted transcription factors. Importantly, the sponge effect is tunable, which we show by changing plasmid copy number and also the sequence of the sponge. As a metabolic engineering application, we use arginine biosynthesis as a showcase and design synthetic sponge systems for the arginine production pathway repressor ArgR. Our results show that metabolic flux to the arginine production pathway is increased by the introduction of a synthetic sponge. Using liquid chromatography – mass spectrometry (LC-MS), we demonstrate that production is increased 16.7-fold compared to the parental production strain without a detectable growth difference. Further, we demonstrate that the sponge-based production strain shows higher genetic stability than a similar system using an *argR* knock-out. After a cyclic culturing process to simulate the potential for genetic drift during scale up, we found that 87% of the knock-out strains obtained mutations, while no mutations were detected in the sponge-based production strain. Lastly, we establish the use of sponge libraries for phenotypic screening. Using a combinatorial library of sponges, we significantly increased pinene tolerance with a dual sponge containing binding sites for regulators of tolerance genes. These results indicate that transcription factor sponges have excellent potential as a tool for metabolic engineers due to their low fitness cost, high genetic stability, and compact size.

## RESULTS

### Altering gene expression with the sponge system

To investigate the effects of the transcription factor sponge, we first employed two widely used inducible promoters, P_lac_ and P_tet_, which are controlled by the transcription factors LacI and TetR, respectively. To construct a sponge system for each, we inserted a LacI or TetR binding site sequence acquired from the P_lac_ or P_tet_ promoter region into a high copy plasmid. To avoid unexpected transcription from insertion of the decoy binding site, we placed a transcription terminator immediately downstream of the sponge sequence (Fig. 1a).

**Figure 1.**
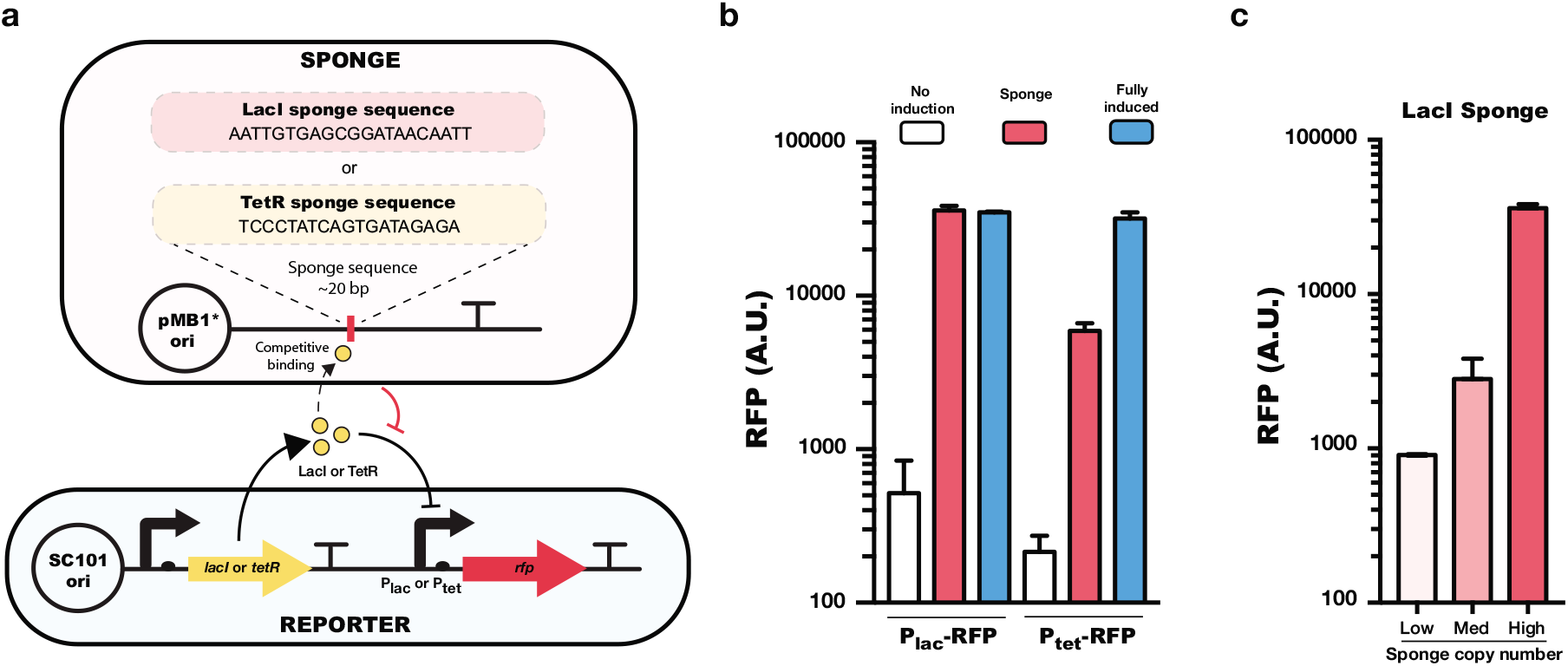
Sponge system for the transcriptional repressors LacI and TetR. **(a)** Schematic view of the sponge plasmid design for LacI and TetR. The LacI or TetR sponge sequence is inserted into a plasmid, where the sequence is immediately followed by a transcription terminator. The copy number of this plasmid depends on the experiment, but pMB1* is shown as an example. The reporter plasmid contains the P_lac_ or P_tet_ promoter driving expression of the gene for red fluorescent protein (*rfp*) on a low-copy SC101 origin plasmid. **(b)** P_lac_ or P_tet_ promoter expression levels with no induction (no sponge, no inducer), with sponge (no inducer) and fully induced (no sponge, 1 mM IPTG, or 100 nM aTc). **(c)** Regulatory effects of the LacI sponge system in low copy (p15A), medium copy (ColE1), or high copy (pMB1*) origin of replication plasmids. Error bars show standard errors from n = 3 biological replicates.

First, to test the effects of the LacI and TetR sponges, we co-transformed the sponge system with a reporter plasmid consisting of either P_lac_ or P_tet_ driving expression of the gene for red fluorescent protein *(rfp)* (Fig. 1a). We found that the sponges can effectively activate by relieving repression of a promoter. The LacI sponge allowed the P_lac_ promoter to be expressed at a level comparable to saturating-levels of IPTG induction. The TetR sponge results were similar, with significantly elevated expression from P_tet_, though in this case the sponge system achieved expression levels that were not quite as high as in the fully induced conditions (Fig. 1b).

To test whether the titration effect is a function of the copy number of the sponge, we introduced the LacI sponge into plasmids with different replication origins: p15A, ColE1, and pMB1*, representing low (~10), medium (~20), and high (~500) copy numbers plasmids. Using the P_lac_-RFP reporter, we observed a clear trend of increased transcriptional activity as the copy number of the sponge plasmid is increased (Fig. 1c). This suggests that the sponge effect can be tuned by controlling copy numbers in the cell.

### Controlling a native metabolic pathway using the synthetic sponge system

We next sought to test whether the sponge effect could be used to control the activity of a native metabolic pathway. To do this, we focused on arginine biosynthesis as a model system. The arginine production pathway is regulated by the transcription factor ArgR, which controls 31 targets (29) and functions as a strong repressor in the presence of arginine (32), preventing its overproduction (Fig. 2a).

**Figure 2.**
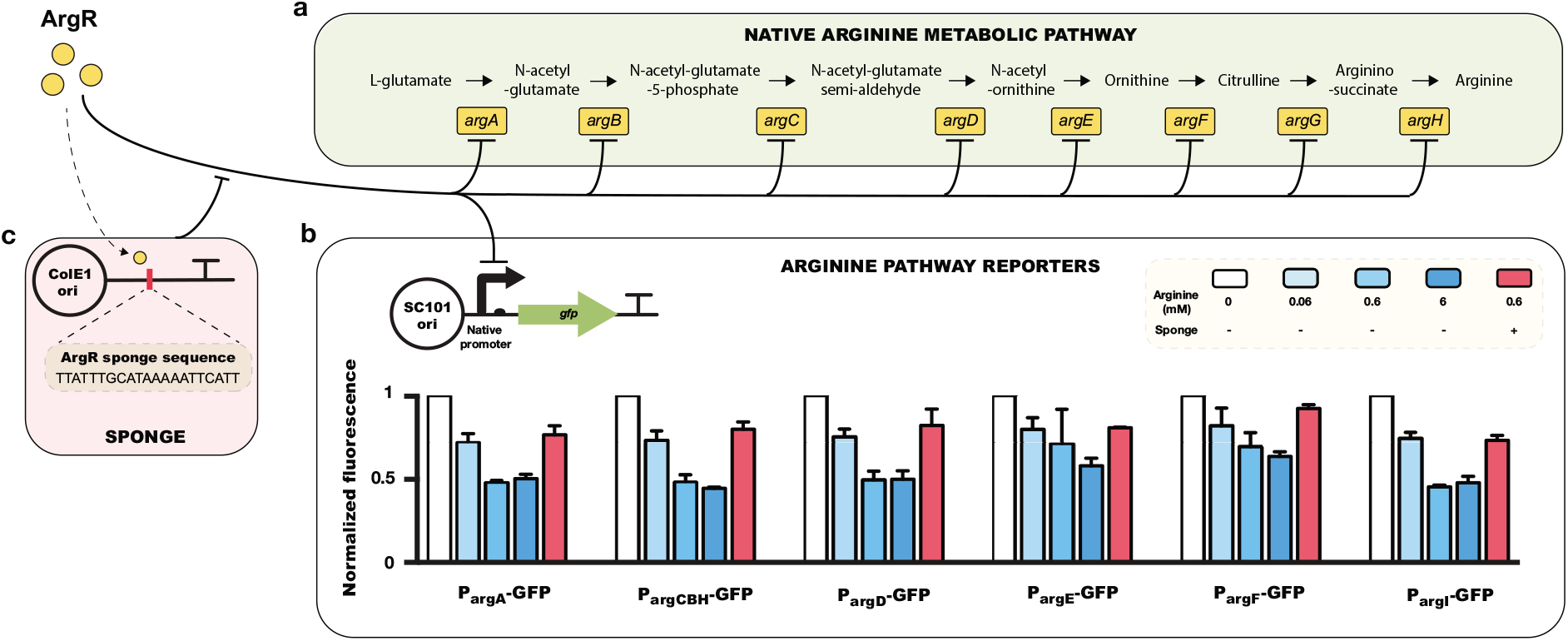
ArgR sponge for regulating the native arginine biosynthesis pathway. **(a)** Schematic view of the native arginine production pathway in *E. coli.* **(b)** Up-regulation of arginine production pathway genes by introduction of the ArgR sponge system in a plasmid with ColE1 origin. Expression of reporters are normalized to the values from the 0 mM arginine case. Error bars show standard error from n = 3 biological replicates. **(c)** Design of the ArgR sponge system.

To establish a baseline for the minimum and maximum levels of ArgR regulated promoter activities, we conducted tests with and without arginine on native promoters driving green fluorescent protein (*gfp*) expression (Fig. 2b). We employed a set of six transcriptional reporters for the arginine production pathway: P_argA_-GFP, P_argCBH_-GFP, P_argD_-GFP, P_argE_-GFP, P_argF_-GFP, and P_argI_-GFP. We measured the transcriptional activity of all six reporters upon supplementation with 0.06 mM, 0.6 mM, or 6 mM arginine. In the presence of arginine, ArgR binds arginine causing repression of genes in the pathway. As expected, increasing concentrations of arginine have a clear negative impact on the transcriptional activity of all reporters (Fig. 2b).

We next constructed a sponge system for ArgR to test whether we could use it to restore transcriptional activity of the production pathway, even in the presence of arginine. To test this, we designed an ArgR sponge, which consists of a 21bp artificial consensus sequence (Fig. 2c). To determine the consensus, we calculated the strict consensus region of all 31 known ArgR binding sites based on its position weight matrix (30). We co-transformed the sponge system with each of the arginine pathway reporters and measured the transcriptional activity in the presence of 0.6 mM arginine. We chose 0.6 mM arginine because it is a production-relevant concentration (7) and most arginine pathway genes reach a repression plateau above this concentration (Fig. 2b). In the strain containing the ArgR sponge with arginine present, we observed a notable restoration of transcriptional activity with all six reporters, suggesting that the sponge system can effectively titrate ArgR away from its genomic targets and up-regulate the arginine production pathway (Fig. 2b).

### Tunable regulation of the transcription factor sponge

To evaluate whether the sponge effect can be tuned by modifying the sequence of the ArgR binding site, we constructed a library of ArgR sites (Fig. 3a) and tested the sponge effect using a P_argA_-GFP reporter. We found that although all members in the ArgR sponge library share the same position weight matrix logo, sequence differences in the non-conservative region of the ArgR binding sites resulted in variants with a range of impacts in 0.6 mM arginine treatment, from no effect to full restoration of transcriptional activity, suggesting that the sponge effect is highly dependent on the DNA sequence and is tunable if the binding sequence is modified (Fig. 3b).

**Figure 3.**
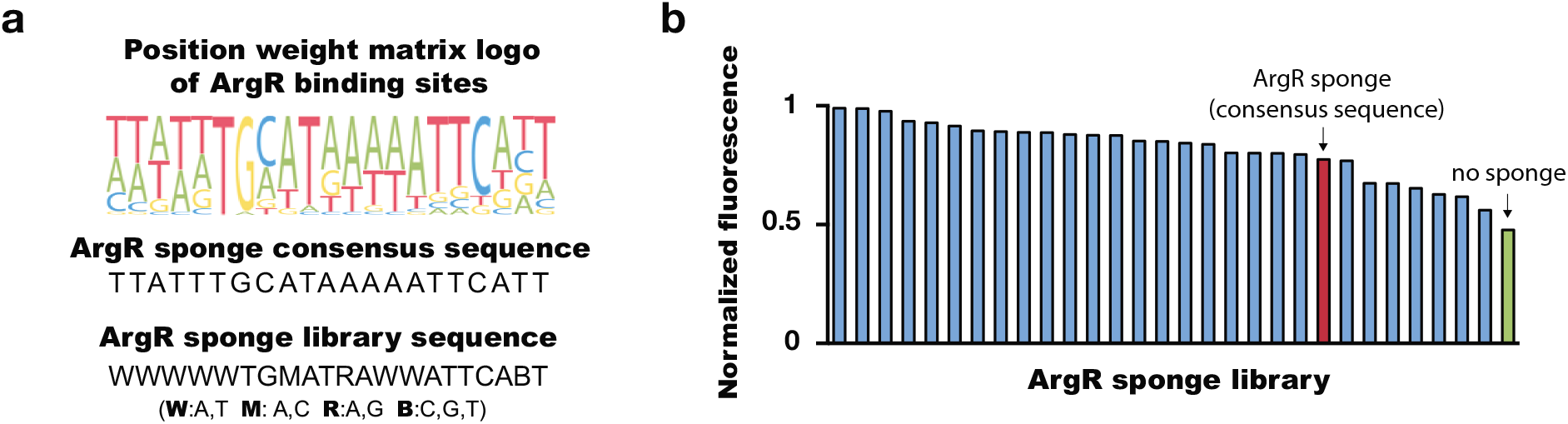
Tunable control of the ArgR sponge system. **(a)** Sequences in the ArgR sponge library. **(b)** Regulatory effects of different sequences in the ArgR sponge library in 0.6 mM arginine treatment. Library members are sorted from high (most effective as a sponge) to low (least effective).

Because the sponge effect is a function of its copy number (Fig. 1c), we reasoned that an alternative way to engineering tunability would be to varying the plasmid copy number. To achieve this, we employed a plasmid with an IPTG-inducible phage P1 replication system for tunable copy number (31) and incorporated the ArgR consensus sponge into the inducible copy number plasmid (Fig. S1a). We measured transcriptional activity of an arginine pathway reporter (P_argA_-GFP) in the presence of arginine. As expected, we observed a trend of increased expression of the arginine pathway reporter with increased IPTG induction in 0.6 mM arginine treatment, suggesting that the sponge effect can be controlled by fine-tuning copy number via exogenous addition of an inducer (Fig. S1b). Further engineering, such as reducing the leaky expression level of RepL to decrease the baseline plasmid number could further improve tunability.

### Enhanced arginine yields without a growth deficit using the synthetic sponge system

To confirm the sponge effect and quantify its ultimate impact on arginine production, we introduced the ArgR sponge into an arginine production strain. This base production strain harbors a plasmid expressing a mutated version of *argA, argA* (H15Y), which we denote *argA*,* where allosteric feedback is removed (Fig. 4a). To quantify arginine production, we used LC-MS to measure the yields of arginine in strains with and without the sponge after 24 hours of fermentation. Since we observed significant up-regulation of arginine production pathway genes with our ArgR sponge system (Fig. 2b), we reasoned that arginine production should also increase with the ArgR sponge. Indeed, we observed a 16.7-fold increase in arginine yields after co-transforming ArgA* with the ArgR sponge system compared to the same strain without the sponge (Fig. 4b). These results confirm that the sponge system can effectively steer metabolic pathway activity, increasing yields. Importantly, even with the increase in arginine production, we observed no detectable growth differences compared with the wide type strain (Fig. 4c). This result is in contrast to strategies based on an *argR* knock-out, which have a significant growth deficit (Fig. 4c). These results suggest that the sponge system is an effective tool for redirecting metabolic flux that imposes a low burden to the cell.

**Figure 4.**
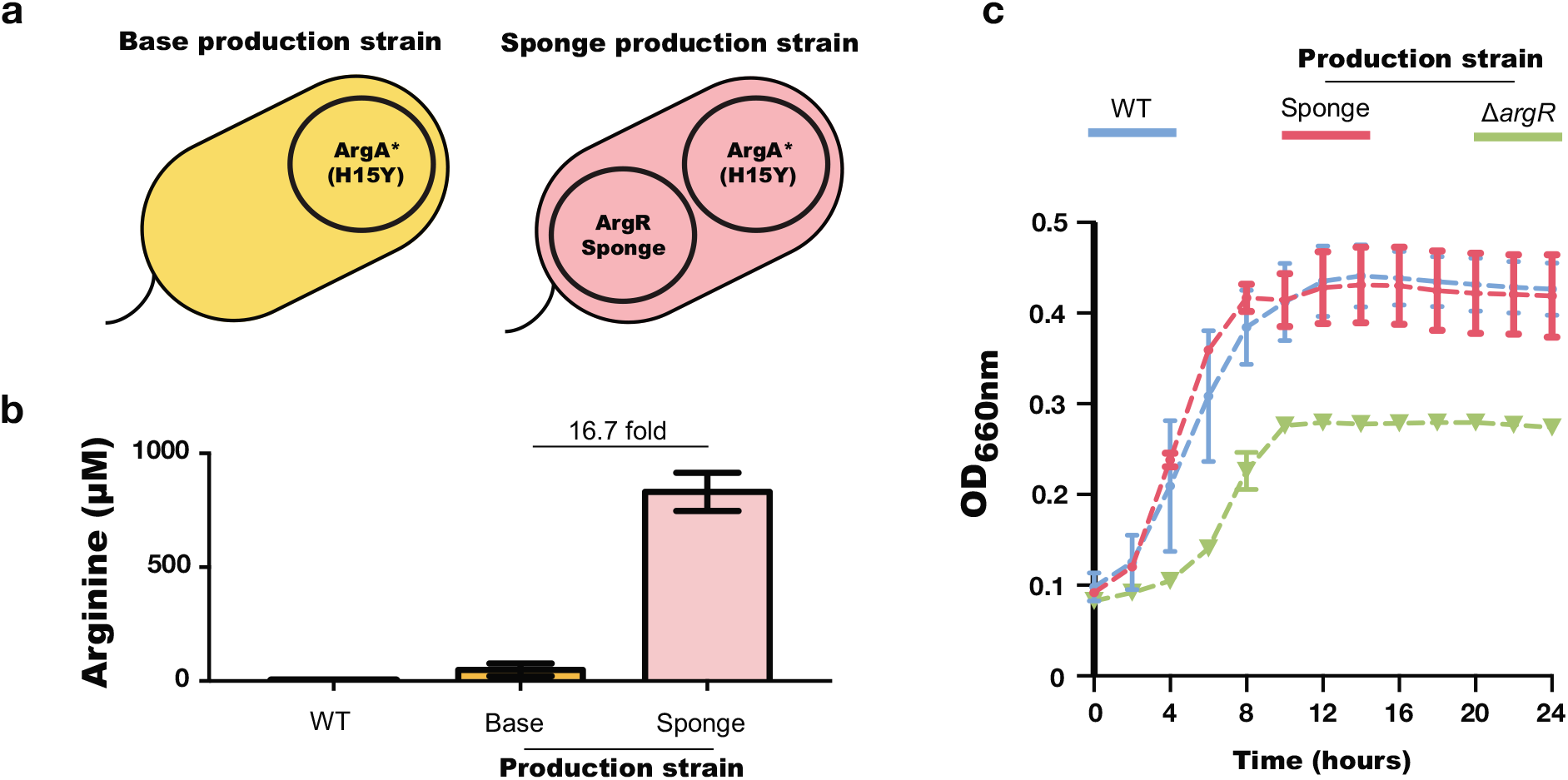
Yield and growth curves of arginine production strains. **(a)** Schematic view of the arginine production strains. **(b)** Arginine yield of production strains measured by LC-MS. **(c)** Growth curves of different arginine production strains. Error bars show standard deviations from n = 3 biological replicates.

### Increased genetic stability using the sponge system

Engineered cells that exhibit a growth deficit can eventually be out-competed by low-productivity counterparts that acquire a mutation that restores fitness. Because the production strain based on the *argR* deletion exhibits a significant growth deficit, we reasoned that production strains based on the knock-out strategy might be mutation prone, since mutations occurring in the ArgA* production plasmid could recover growth, but decrease yields. However, since the sponge-based production strain shows negligible fitness differences compared to wild type, we anticipated that the propensity for mutation would be lower.

To compare the genetic stability between *argR* deletion-based and sponge-based production strains, we quantified the mutations occurring during continuous culture by sequencing the ArgA* plasmid and sponge plasmid at the end of each cycle (Fig. 5a). In the knock-out production strain (Δ*argR*), we found that cells containing plasmids with mutations within the *argA\** coding sequence quickly took over the culture; 7 out of 8 colonies that we sequenced were mutated by cycle 6 (Fig. 5b). Of these mutated plasmids, 6 of the 7 contained Y15H, which reverts *argA\** to *argA* by restoring allosteric feedback (Table S1). In contrast, the sponge-based production strain showed high genetic stability, and the colonies that we sequenced contained no mutations in either *argA\** or the sponge region of the production plasmid or the sponge plasmid itself (Fig. 5b). This result suggests that the sponge-based production strategy has higher genetic stability than the burdensome *argR* deletion-based version, allowing for the maintenance of intact production from heterologous elements after many cycles of cell division.

**Figure 5.**
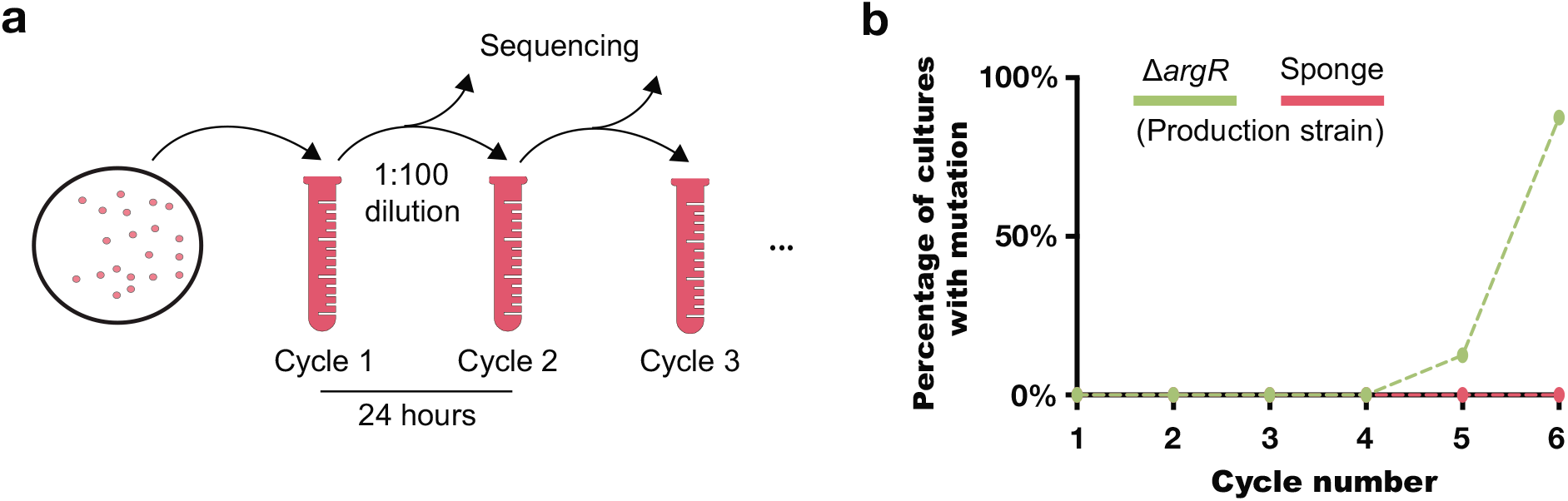
Genetic stability of arginine production strains. **(a)** Schematic diagram of genetic stability test. Eight different bacterial colonies from each production strain were subjected to continuous culture. Every 24 hours, we isolated and sequenced regions of interest from the bacterial cultures. The target sequencing regions of interest were *argA\** for the *ΔargR* production strain, and *argA\** and the sponge region in the sponge-based production strain. **(b)** Percentages show the number of bacterial cultures with a mutation out of eight sequenced.

### Improved pinene tolerance using a multiplexed sponge library

Since transcription factor sponges have the ability to perturb transcriptional programs and alter phenotypes, we reasoned that they could also enhance tolerance phenotypes. A library approach to tolerance screening is easily adaptable to sponge systems since the binding sequences are short and can be created by annealing oligos. Moreover, sponges do not rely on transcription or translation, making them amenable to high efficiency multi-part Golden Gate assemblies (33) without the need to ensure in-frame inserts. By matching oligo overhang sequences, or using palindromic overhang sequences, libraries of multiple sponge inserts can be created in streamlined one-pot reactions. As a showcase of sponge tolerance libraries, we sought to enhance tolerance to α-pinene, an important monoterpene that can be produced in *E. coli* (34). Pinene is of interest to metabolic engineers as it has many potential uses, such as an alternative jet fuel, flavoring and fragrance additive, and a therapeutic agent (34–36). Production of pinene is toxic to *E. coli* and growth is inhibited in 0.5% pinene (v/v). Several studies have demonstrated that *E. coli* can cope with pinene-induced stress using endogenous genes (37–39).

To test whether we could rapidly screen for improved pinene tolerance using a multiplexed library approach, we created a dual sponge combinatorial library. The library is based on regulators of genes known to play a role in pinene tolerance (Table S2). It was constructed in a pooled, single-pot reaction, highlighting the simplicity of the sponge-based design (Fig. 6a). Single colonies were subjected to 0.5% pinene and we monitored growth to identify tolerant variants (Fig. 6b). Several colonies showed increased tolerance relative to the wild type strain. We sequencing the sponge insert and identified unique pairs of binding sites in the top tolerance strains (Table S3). In order of highest endpoint optical density, the sponge pairs were SoxR-UlaR, AcrR-AcrR, and SoxR-MarR. SoxR regulates dozens of downstream genes involved in stress response (40). UlaR is a regulator of the *ula* operon that is responsible for L-ascorbate metabolism. UlaE has been reported to significantly enhance pinene tolerance (39), however, the specific mechanism by which this operon increases tolerance is not well understood. AcrR regulates expression of the multidrug efflux pump AcrAB-TolC, which is also known to increase pinene tolerance (37). These results demonstrate the straightforward application of multiplexed sponge-based approaches to library selection.

**Figure 6.**
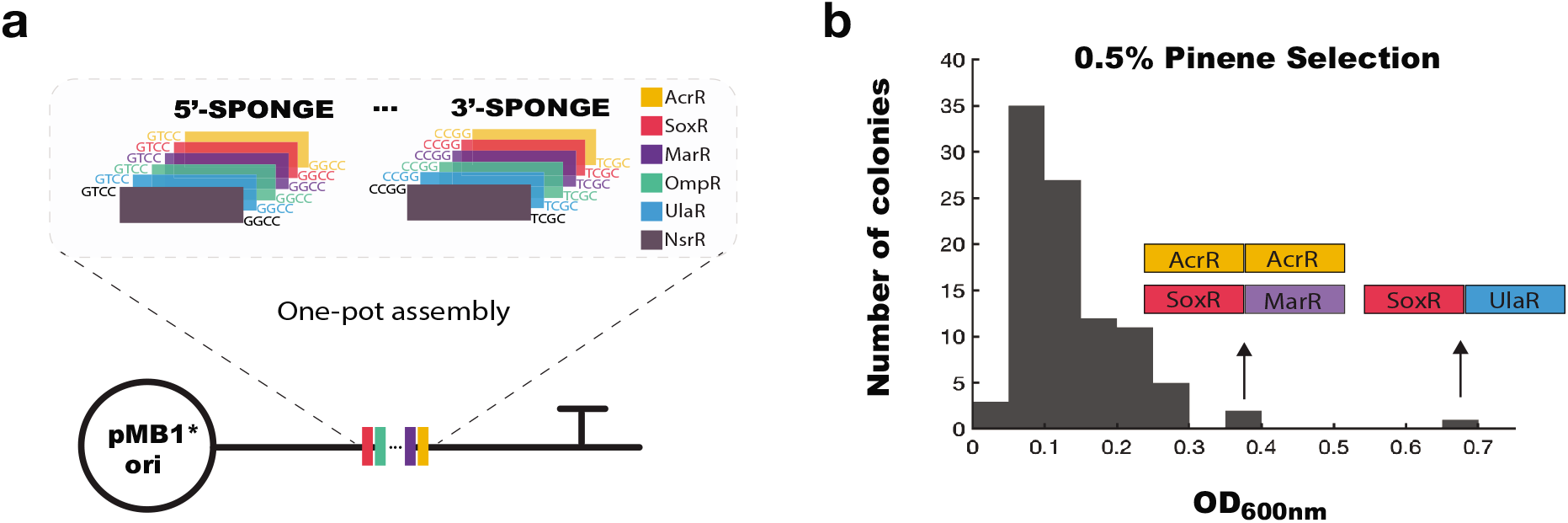
Multiplexed sponge library for pinene tolerance. **(a)** Cloning scheme to create combinatorial sponge libraries. The transcription factor binding site sequences are on oligos that are annealed, followed by a single-pot Golden Gate assembly. **(b)** Optical density at 600nm (OD600) of colonies grown in 0.5% pinene (v/v) for 12 hours. Sponge sequences of the top three sponge combinations are shown.

## DISCUSSION

We have harnessed decoy binding sites to titrate transcription factors in order to regulate expression of genes without reducing fitness, therefore increasing yield and genetic stability in a metabolic engineering context. We have shown that transcription factor sponges are an effective tool for altering gene expression for both native and heterologous targets. Importantly, the effect of the sponge can be tuned by changing its copy number or DNA sequence. As an application, we used a sponge system to control arginine biosynthesis and showed that it can regulate metabolic flux by increasing transcriptional activity of the arginine production pathway, resulting in a more than 16-fold increase in arginine production compared to a parental strain lacking the sponge. In contrast to production strains based on an *argR* knock-out, the sponge system exhibits no detectable growth difference compared to wild type. This suggests that using the sponge to selectively titrate away transcription factors may have a much smaller burden compared to alternative strategies. Since fitness deficits can compromise genetic stability, we also compared the number of mutations between alternative designs. We found that the production strain based on the sponge system has higher genetic stability than the knock-out system. Further, by screening for pinene tolerance, we have shown that the method is highly amenable to multiplexing. It is feasible to scale up library diversity by increasing the number of sponge inserts by using palindromic overhangs or increasing the number of sponges in the library.

The sponge approach is a powerful system for selectively blocking the effects of transcription factors and regulating metabolic pathways in bacteria. When applied to arginine production, the sponge approach increased yield without a detectable growth deficit and exhibited high genetic stability. We also demonstrated tolerance enhancing sponge effects, which can further increase productivity and stability of engineered strains. Our results suggest that using the sponge system in metabolic engineering applications can allow researchers to develop strains that are productive while maintaining favorable genetic stability and growth characteristics.

The design simplicity adds to the appeal of sponged based transcriptional regulation and opens this method up for many future applications. For example, this approach can extend to organisms beyond *E. coli* where tunable expression systems and synthetic biology tools are more limited. Prokaryotic systems that rely heavily on negative regulation would likely have the ability to be regulated by transcription factor sponges. In metabolic engineering, this is of particular interest for non-model organisms that have the ability to grow on desired feedstocks, such as cellulosic biomass or even through photosynthesis. In these non-model organisms, sponges could potentially be applied to steer biosynthesis towards desired end products.

## MATERIAL AND METHODS

### Strains

We used *E. coli* BW25113 as the wild type strain. *ΔargR* was derived from *E. coli* BW25113 and we deleted the *argR* gene using homologous recombination (27). We used the forward primer 5’-AAG CAA GAA GAA CTA GTT AAA GCA TTT AAA GCA TTA CTT AAA GAA GAG AAg tgt agg ctg gag ctg ctt c-3’ and the reverse primer 5’-CCT GGT CGA ACA GCT CTA AAA TCG CTT CGT ACA GGT CTT TGA CTG TGA AAa ttc cgg gga tcc gtc gac c-3’. Capitalized letters indicate the homologous recombination extension.

The ArgA* production strain was created by transforming *E. coli* BW25113 with an ArgA* production plasmid containing *P_argA_-argA* (H15Y). To construct the ArgA* production plasmid, we amplified the promoter and gene region of *argA*, P_argA_-*argA*, from *E. coli* BW25113 using the forward primer 5’-GCCTCTCCCGAGCAAAAG −3’ and reverse primer 5’-TTACCCTAAATCCGCCATCAAC −3’. We then introduced mutations in the PCR product of P_argA_-*argA* to create P_argA_-*argA* (H15Y) using the forward primer 5’-GGG ATT CCG CTA TTC AGT TCC −3’ and reverse primer 5’-TTA CCC TAA ATC CGC CAT CAA C −3’. P_argA_-*argA* (H15Y) was then cloned on the low-copy (SC101) plasmid pBbS5C from Ref. (28).

### Transcription factor sponge plasmid construction

#### LacI sponge plasmid

We cloned the LacI binding site sequence AAT TGT GAG CGG ATA ACA ATT into the p15A replication origin plasmid pBbA5A, or ColE1 replication origin plasmid pBbE5A, using the forward primer 5’-AAT TGT GAG CGG ATA ACA ATT cca tcg ttg aac agt acg aac −3’, and reverse primer 5’-AAT TGT TAT CCG CTC ACA ATT cca tca aac agg att ttc gcc −3’. For cloning into the pMB1 (high copy derivative) plasmid pUC19, we used the forward primer 5’-AAT TGT GAG CGG ATA ACA ATT taa tgc agc tgg cac gac −3’, and reverse primer 5’-AAT TGT TAT CCG CTC ACA ATT ggt ttg cgt att ggg cgc −3’. Capitalized letters indicate the homologous region. All plasmids were derived from the BioBrick library described in Lee *et al.* (28).

#### TetR sponge plasmid

We used the forward primer 5’-TCC CTA TCA GTG ATA GAG Ata atg cag ctg gca cga c −3’, and reverse primer 5’-TCT CTA TCA CTG ATA GGG Agg ttt gcg tat tgg gcg c −3’ to clone the TetR binding site sequence TCCCTATCAGTGATAGAGA into the pMB1 (high copy derivative) plasmid pUC19.

#### ArgR consensus sponge plasmid

For the ArgR consensus sponge plasmid, we introduced the consensus ArgR binding site sequence TTA TTT GCA TAA AAA TTC ATT into the ColE1 replication origin plasmid pBbE5A using the forward primer 5’-TTA TTT GCA TAA AAA TTC ATT TGT ATG CAC Agc tga agg tcg tca ctc ca −3’, and reverse primer 5’-AAT GAA TTT TTA TGC AAA TAA CAG TCA GCC CCc cac cgt ctt tca gtt tca ga −3’.

#### ArgR sponge library plasmid

For the ArgR sponge library plasmid, we introduced the ArgR binding site sequence with randomized sequence WWW WWT GMA TRA WWA TTC ABT (W:AT; M:A,C; R:A,G; B:C,G,T) into the ColE1 replication origin plasmid pBbE5A. The position weight matrix was calculated using the 31 ArgR binding sites listed in Ref. (29) using the method described in Ref. (30).

#### ArgR sponge inducible copy number plasmid

For the ArgR consensus sponge plasmid, we introduced the consensus ArgR binding site sequence TTA TTT GCA TAA AAA TTC ATT into a plasmid system with an IPTG-inducible phage P1 replication system with tunable copy number (31).

### LacI and TetR sponge test

Bacteria were cultured in M9 minimal medium with 5 g/L glucose at 37°C with 200 rpm shaking. Overnight cultures inoculated from a single colony were diluted 1:50 in M9 minimal medium with 5 g/L glucose and selective antibiotics for plasmid maintenance, where required. The diluted cultures were then precultured for 3h, 1mM IPTG or 100nM aTc was added (where indicated), then grown at 37 °C with 200 rpm shaking. Red fluorescent protein (RFP) readings (excitation 580 nm, emission 610 nm) were taken using a BioTek Synergy H1m plate reader (BioTek, Winooski, VT) after 18 hours of incubation at 37 °C with 200 rpm shaking.

### ArgR sponge test

Bacteria were cultured in M9 minimal medium with 5 g/L glucose at 37°C with 200 rpm shaking. Overnight cultures inoculated from a single colony were diluted 1:50 in M9 minimal medium with 5 g/L glucose and selective antibiotics for plasmid maintenance, where required. The diluted cultures were then precultured for 2h, arginine and IPTG were added when needed at the concentrations indicated in figure captions, then cultures were grown at 37 °C with 200 rpm shaking. Green fluorescent protein (GFP) readings (excitation 480 nm, emission 510 nm) were taken using a BioTek Synergy H1m plate reader after 4 hours of incubation with shaking at 37 °C.

Normalized fluorescence was calculated with the following equation: 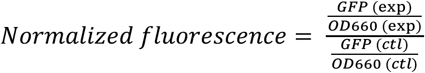, where GFP (exp) and OD660 (exp) are GFP readings and optical density readings of arginine pathway reporters in the strain with or without the ArgR sponge and with or without arginine treatment. GFP (ctl) and OD660 (ctl) are GFP readings and optical density reading for the strain harboring the arginine pathway reporters without arginine treatment.

### Arginine production experiments

Overnight cultures of the production strains were diluted 1:50 in 5 ml M9 minimal medium with 5 g/L glucose with antibiotics, when appropriate. Diluted cultures were then cultured for 24h at 37°C with 200 rpm shaking. Bacterial cultures were then placed on ice and lysed with 10 cycles of sonication (10 seconds ON, 30 seconds OFF, 20% amplitude). 40 μl of cell lysate was mixed with acetone at a 1:8 ratio, vortexed and kept on ice for 30 minutes. Samples were then centrifuged at 15,000 rcf for 10 minutes at 10°C to pellet proteins and lipids. Supernatant was transferred to a new tube, leaving behind the protein pellet. The sample was then dried in a speed vacuum centrifuge and then reconstituted by adding 40μl H20 with 0.1% formic acid and vortexed. Samples were placed on ice for 15 minutes and centrifuged to remove any residual protein or lipid. Supernatant was then transferred to a glass LC vial with a glass insert and measured using an Agilent HPLC 1100 series auto sampler.

### LC-MS/MS

An Agilent HPLC 1100 series was used with a Phenomenex Kinetex 2.6μm F5 100Å 150mm × 2.1mm column (PN 00F-4723-AN). Arginine was analyzed on a Sciex API 4000 mass spectrometer triple quadrupole in positive polarity with a targeted Q1 Mass of 175.100 Da and a Q3 mass of 70.000 Da, with a dwell time 20.0ms, declustering potential (DP) 50.0 volts, entrance potential (EP) 10.0 volts, collision energy (CE) 32.0 volts, and a collision cell exit potential (CXP) 9.0 volts. Sample was injected at 5 μL onto the column and was analyzed at a gradient of 97% H_2_O and 0.1% formic acid (Buffer A) and 3% acetonitrile and 0.1% formic acid (Buffer B) at 0-2.0min, 3% Buffer B at 2.0min to 95% Buffer B at 7 min, 95% Buffer B to 8.0min, to 3.0% Buffer B at 8.5-10.0 min at a flow rate of 200 μl/min. Sciex MultiQuant 3.0.3 software was used to analyze data and calculate concentrations from a linear plot of arginine standards.

### Genetic stability test

ArgA*/Δ*argR* or ArgA*/sponge strains were cultured in M9 minimal medium with 5 g/L glucose at 37°C with 200 rpm shaking. Overnight cultures inoculated from a single colony were diluted 1:50 in M9 minimal medium with 5 g/L glucose and antibiotics for plasmid maintenance and cultured for 24 hours. We refer to this as cycle 1. Every 24 hours, we diluted the culture 1:100 in fresh M9 minimal medium with 5 g/L glucose and antibiotics and repeated this procedure until cycle 6.

Before dilution each cycle, we isolated DNA to sequence regions of interest. The target regions of interest for sequencing were the coding sequence for *argA*\* for the Δ*argR* production strain, and *argA*\* and the sponge region in the sponge-based production strain. We isolated plasmid from the bacterial cultures using the GenCatch™ Plasmid DNA Mini-Prep Kit. Isolated plasmids were then amplified with PCR with the following primers and sequenced using the forward PCR primers. For the ArgA* plasmid we used forward primer 5’-GCC TCT CCC GAG CAA AAG −3’ and reverse primer 5’-TAT AAA CGC AGA AAG GCC CAC −3’. For the sponge plasmid, we used forward primer 5’-CTG CGT GGT ACC AAC TTC C −3’ and reverse primer 5’-CCG AAC GCC CTA GGT ATA AAC −3’.

### One-pot sponge library

To construct a dual sponge library, forward and reverse oligos containing each transcription factor sponge site with 5’ overhangs were annealed by heating the oligo mix (1 μM) to 95 °C and allowing the heat block to return to room temperature. 1 pmol of the oligo mix was added to a 20 μl Golden Gate reaction containing 100 ng destination plasmid (pATT-DEST, Addgene #79770), 10 units BsaI (NEB #R3733), 10 units PNK (Thermo Fisher), and 1 unit T4 ligase with associated buffer (Promega). The following thermocycler program was run: 25 cycles of 37°C for 2min and 16°C for 5min, followed by 60°C for 10min (final digestion) and 80°C for 10min (heat inactivation). Successful sponges replace *lacZ* on the plasmid.

### Pinene tolerance screen

A combinatorial dual sponge library was created to screen for pinene tolerance. Colonies from the transformed Golden Gate reaction were picked and added to individual wells of a 96-well plate containing LB with 0.5% α-pinene (v/v). After 12 hours, the OD600 of the plate was measured and the cultures with the highest growth were sequenced to identify the incorporated sponges.

## ACKNOWLEDGEMENT

We thank Sarah Nemsick for early work on transcription factor sponge construction and screening; Dr. Nadia Sampaio assistance with sponge evolution experiments; Dr. Nicholas Rossi, Dr. Nadia Sampaio, Dr. Jean-Baptiste Lugagne, and Ariel Langevin provided input during manuscript development.

