## Supplementary data for "Programmable gene regulation for metabolic engineering using decoy transcription factor binding sites"

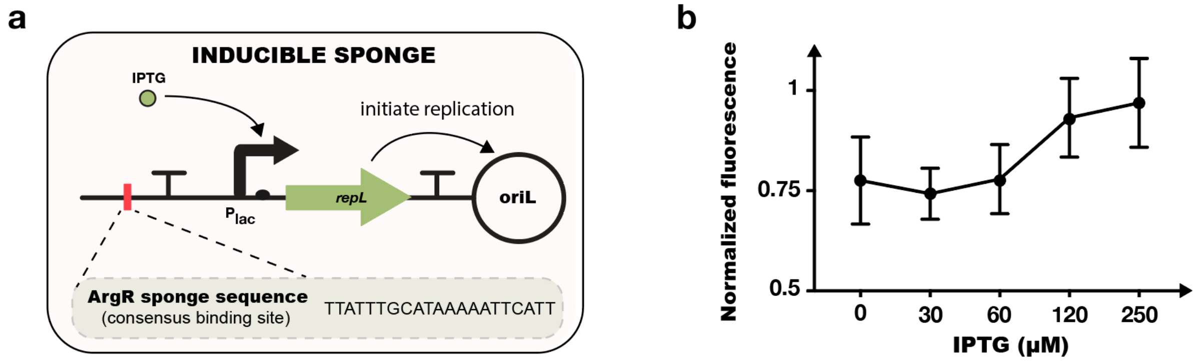


**Figure S1. Tunable control of the ArgR sponge system with inducible copy number plasmid. (a)** Schematic view of the design of the inducible ArgR sponge system. **(b)** ArgR sponge effect is enhanced as IPTG increases the plasmid copy number (0.6 mM arginine treatment). Error bars show standard error from n = 3 biological replicates.

**Table S1. Mutations found in ArgA* in the strain of ArgA*/Δ*argR* at cycle 6**

| Colony # | Mutation |
|---|---|
| 1 | *A197R* |
| 2 | None |
| 3 | *Y15H* |
| 4 | *Y15H* |
| 5 | *Y15H* |
| 6 | *Y15H* |
| 7 | *Y15H, T93H* |
| 8 | *Y15H* |

**Table S2. Pinene tolerance library sponge sequences**

| Sponge | Forward Sequence |
|---|---|
| AcrR | GATTTACATACATTTNTGAATGTATGTA |
| SoxR | GAACCCTCAAGTTAACTTGAGG |
| MarR | ACTAATTACTTGCCAGGGCAAGTAAT |
| OmpR | TTTACTTTTGGTTACATCTA |
| UlaR | TGATTAATCATGAACAATCA |
| NsrR | GATGCATTTAAAATACATC |

**Table S3. Top five hits with associated $OD_{600}$ in pinene tolerance screen**

| Dual Sponge | Colony $OD_{600}$ |
|---|---|
| SoxR-UlaR | 0.694 |
| AcrR-AcrR | 0.389 |
| SoxR-MarR | 0.364 |
| SoxR-OmpR | 0.29 |
| MarR-NsrR | 0.264 |
